# CRISPR/Cas9-APEX-mediated proximity labeling enables discovery of proteins associated with a predefined genomic locus in living cells

**DOI:** 10.1101/159517

**Authors:** Samuel A Myers, Jason Wright, Feng Zhang, Steven A Carr

## Abstract

The activation or repression of a gene’s expression is primarily controlled by changes in the proteins that occupy its regulatory elements. The most common method to identify proteins associated with genomic loci is chromatin immunoprecipitation (ChIP). While having greatly advanced our understanding of gene expression regulation, ChIP requires specific, high quality, IP-competent antibodies against nominated proteins, which can limit its utility and scope for discovery. Thus, a method able to discover and identify proteins associated with a particular genomic locus within the native cellular context would be extremely valuable. Here, we present a novel technology combining recent advances in chemical biology, genome targeting, and quantitative mass spectrometry to develop genomic locus proteomics, a method able to identify proteins which occupy a specific genomic locus.

## Body

Transcriptional regulation is a highly-coordinated process largely controlled by changes in protein occupancy at regulatory elements of the modulated genes. Chromatin immunoprecipitation (ChIP) followed by quantitative polymerase chain reaction (-qPCR), microarrays (-chip), or massively parallel next-generation sequencing (-seq) has been invaluable for our understanding of transcriptional regulation and chromatin structure, both at the individual locus and genome-wide levels (*1*–*6*). However, because ChIP requires the use of antibodies, its utility can often be limited by the presupposition of a suspected protein’s occupancy, and lack of highly specific and high affinity reagents. Previously developed “reverse ChIP” type methods suffer from several drawbacks including loss of cellular and/or chromatin context, extensive engineering and locus disruption, reliance on repetitive DNA sequences, and the need for chemical crosslinking, which reduces sensitivity for mass spectrometric-based approaches (*7*–*11*). Therefore, we sought to develop a method to identify proteins associated with a specific, non-repetitive genomic locus in the native cellular context without the need for crosslinking or genomic alterations. Here, we utilized recent advances in sequence-specific DNA targeting and affinity labeling in cells to develop genomic locus proteomics (GLoPro) to characterize proteins associated with a particular genomic locus.

We fused the catalytically dead RNA-guided nuclease Cas9 (dCas9) (*12*, *13*) to the engineered ascorbate peroxidase APEX2 (*14*) to biotinylate proteins proximal to defined genomic loci for subsequent enrichment and identification by liquid chromatography-mass spectrometry (LC-MS/MS) (Figure 1 A-B). For this proof-of-principle study, dCAS9 was chosen over transcription activator-like effectors (TALEs) or engineered zinc finger nucleases (ZFNs) due to its easily reprogrammable nature of the RNA base pairing to the target locus (*15*). APEX2, in the presence of hydrogen peroxide, oxidizes the phenol moiety of biotin-phenol compounds to phenoxyl radicals that react with surface exposed tyrosine residues, labeling nearby proteins with biotin derivatives (*14*, *16*, *17*). Affinity labeling in cells also circumvents the need for chemical crosslinking, a method used to stabilize biomolecular interactions that severely diminishes LC-MS/MS sensitivity. APEX2 was used instead of other promiscuous biotin ligases due its smaller labeling radius and shorter labeling times (*18*–*20*). The dCas9-APEX2 (*Caspex*) gene was cloned in frame with the self-cleaving T2A peptide and green fluorescent protein (*Gfp*) under the control of a tetracycline response element into a puromycin-selectable piggybac plasmid (*21*) (Figure 1C).

**Figure 1.**
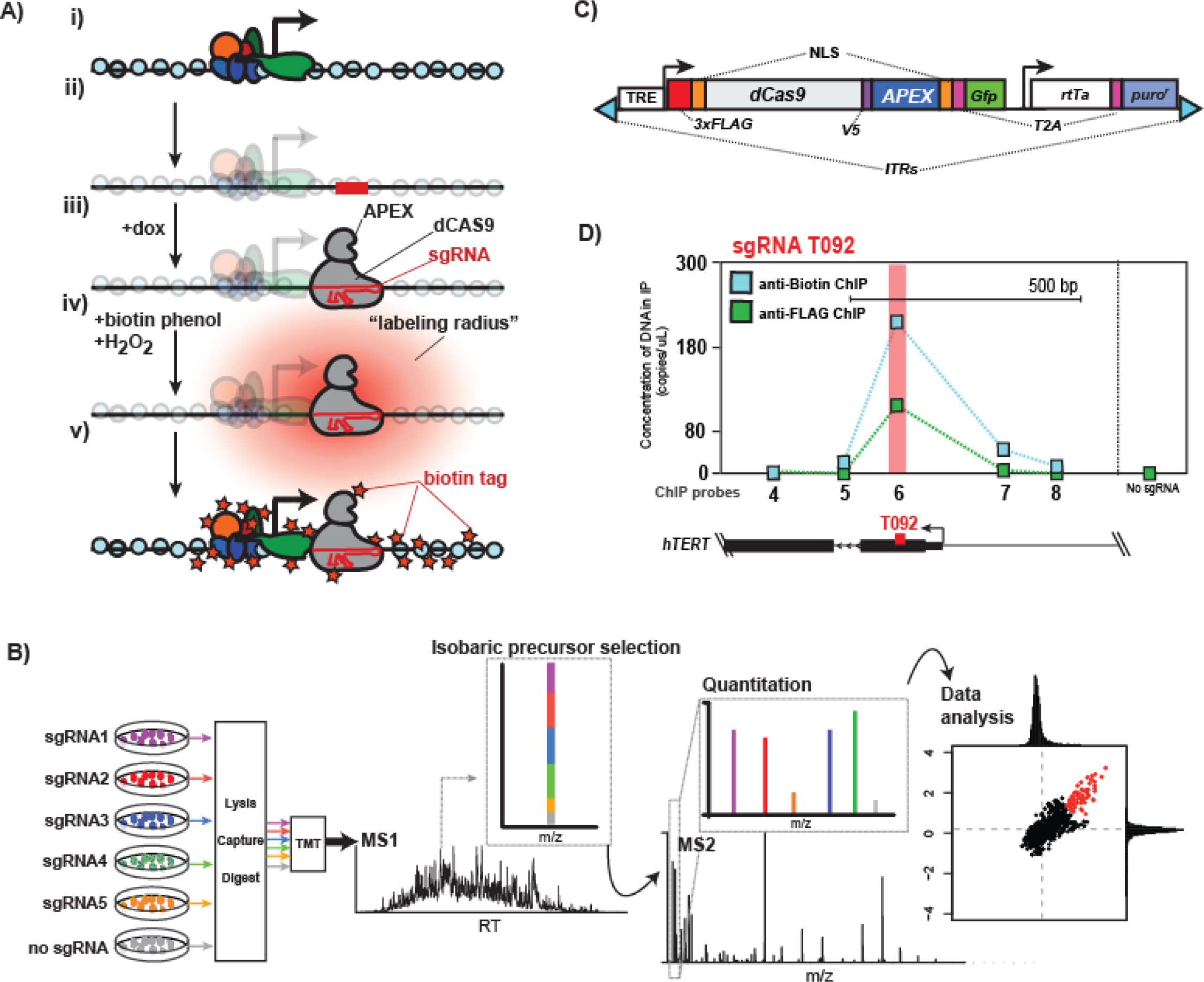
Diagram for Genomic Locus Proteomics workflow. **A)** Illustration of CASPEX targeting and affinity labeling reaction. **i)** A genomic locus of interest is identified. **ii)** A targeting sequence for the sgRNA is designed (red bar). **iii)** CASPEX expression is induced with doxycycline and, after association with sgRNA, binds region of interest. **iv)** After biotin-phenol incubation, H_2_O_2_ induces the CASPEX-mediated labeling of proximal proteins, where the “labeling radius” of the reactive biotin-phenol is represented by the red cloud. **v)** Proteins proximal to CASPEX are labeled with biotin (orange star) for subsequent enrichment. **B)** Workflow for the proteomic aspect of GLoPro. Each individual sgRNA-293T-Caspex line is independently affinity labeled, lysed, enriched for biotinylated proteins by streptavidin-coated beads, digested, and TMT labeled. After mixing, the peptides are analyzed by LC-MS/MS, where the isobarically-labeled peptides from each condition is co-isolated (MS1), cofragmented for peptide sequencing (MS2), and the relative quantitation of the TMT reporter ions are measured. Subsequent data analysis compares the TMT reporter ions for each sgRNA line to the non-spatially constrained no guide control line (grey) to identify reproducibly enriched proteins. **C)** Diagram of Caspex plasmid. NLS, nuclear localization sequence; 3xFLAG, triple FLAG epitope tag; V5, V5 epitope tag; T2A, T2A self-cleaving peptide; GFP, Green fluorescent protein; TRE, Tetracycline response element; rtTA, reverse tetracycline-controlled transactivator; puro^r^; puroMYCin acetyltransferase, ITRs, inverted terminal repeats. **D)** ChIP-qPCR against biotin (blue boxes) and FLAG (green boxes) in 293T-CasPEX cells expressing either no sgRNA (far right) or T092 sgRNA. ChIP probes refer to regions amplified and detected by qPCR as in Supplemental Figure 1. *hTERT* is below to show the gene structure with respect to the sgRNA target (red box).

HEK293T cells were transfected with the *Caspex* plasmid and after 14 days of puromycin selection, single colonies were expanded and characterized for doxycycline (dox) inducible expression of GFP. This clonal, stable line is subsequently referred to as 293T-Caspex. To test whether the CASPEX protein would correctly localize to a genomic site of interest, we expressed a single guide RNA (sgRNA) targeting 92 base pairs (bp) 3’ of the transcription start site (TSS) of the *TERT* gene (sgRNA position is referred to as T092). We chose to focus on the *TERT* promoter (*hTERT*) as *TERT* expression is a hallmark of cancer and recurrent promoter mutations in *hTERT* have been shown to re-activate *TERT* expression (*22*). Biotinylation in T092 sgRNA expressing 293T-CasPEX cells was accomplished by incubating cells with dox for 18 hours, followed by incubation with biotin-phenol for 30 minutes, and finally with hydrogen peroxide for 60 seconds. We then performed ChIP against both the FLAG epitope of CASPEX, and biotin, followed by quantitative PCR (qPCR) of probes tiling *hTERT* (Figure 1D). ChIP-qPCR showed proper localization of CASPEX with the peak of the anti-FLAG signal overlapping with the destination of the sgRNA. The anti-biotin ChIP-qPCR signal showed a similar trend of enrichment, indicating that CASPEX biotinylates proteins in cells within approximately 400 base pairs on either side of its target locus. No enrichment was observed at T092 for the no sgRNA control, which is not spatially constrained to the targeted locus (Figure 1D).

We then created four additional sgRNA constructs tiling *hTERT*: 430T, 107T, T266 and T959, where the number indicates the targeted position relative to *TERT’s* TSS denoted by “T” (Figure 2A). To test whether these sgRNAs were targeted properly, we created 293T-Caspex cells stably expressing the individual sgRNAs. After performing the labeling reaction, these lines were analyzed by ChIP-qPCR against FLAG and biotin. All constructs correctly targeted and labeled the region of interest, showing the peak of enrichment at the sgRNA site (Supplemental Figure 1). While biotinylation was dependent on CASPEX expression, no difference in biotin patterns between *hTERT* sgRNA lines could be seen by Western blot (Supplemental Figure 2). These data demonstrate that CASPEX targeting can be reprogrammed by substitution of the sgRNAs and that proximal protein biotinylation is CASPEX mediated.

**Figure 2.**
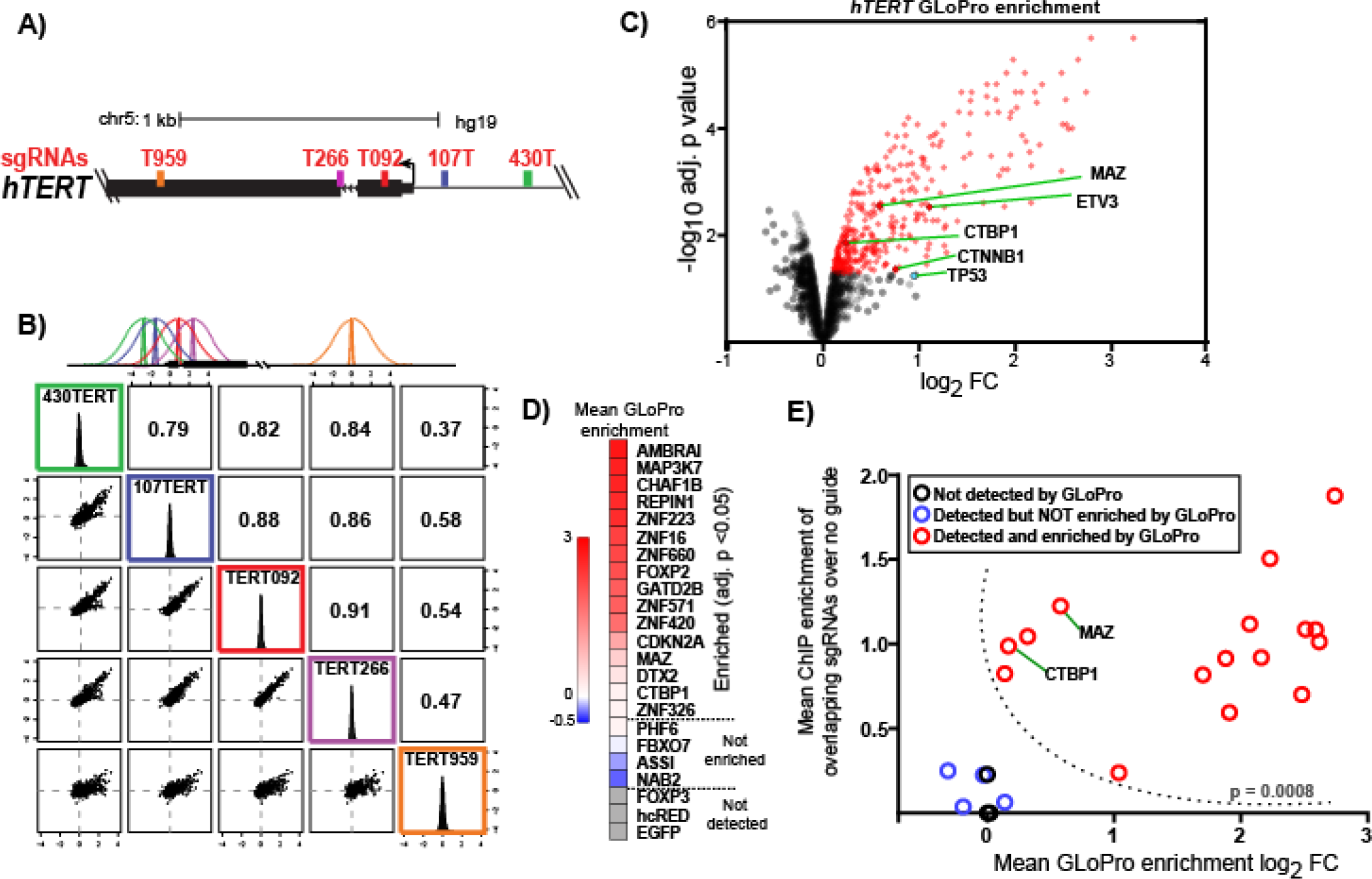
Genomic locus proteomic analysis of *hTERT*. **A)** UCSC Genome Browser representation of *hTERT* (hg19). sgRNAs (colored bars) are shown to scale relative to the transcription start site (black arrow). **B)** Multiscatter plots and Pearson correlation coefficients of log2 fold enrichment values of proteins identified and quantified between hTERT-293T-Caspex cells compared to the no sgRNA control line. **C)** Volcano plot of proteins quantified across the four overlapping *hTERT*-293T-Caspex cell lines compared to the no sgRNA control. Data points representing proteins enriched with an adjusted p-value of less than 0.05 are labeled in red. Proteins known to associate with *hTERT* and identified as enriched by GLoPro are highlighted. TP53, a known *hTERT* binder, had an adj. p val. = 0.058 and is highlighted blue. **D)** Mean GLoPro enrichment values for V5-tagged ORFs selected to ChIP-qPCR corroboration. Red indicates the protein was enriched at *hTERT*, blue that the protein was detected in the analysis but not statistically enriched. Grey proteins were not detected. E) Correlation between ChIP-qPCR and GLoPro enrichment of the four overlapping sgRNAs at *hTERT.* Black, open circles indicate that the protein was not identified by GLoPro. Blue, open circles indicate the protein was identified but was not statistically enriched. Red open circles indicate proteins that are enriched according to the GLoPro analysis. Previously described *hTERT* binders are labeled. Dotted line separates ChIP-qPCR data tested for statistical significance via the Mann-Whitney test, and the p-value is shown.

To test whether CASPEX could identify proteins associated with *hTERT*, we enriched biotinylated proteins with streptavidin-coated beads from *hTERT* -targeted 293T-Caspex lines, followed by analysis with quantitative LC-MS/MS. Biotinylation was initiated in the five individual *hTERT* targeting 293T-Caspex lines 18 hours after addition of doxycycline, along with the no guide control 293T-Caspex line. Whole cell lysates from each individual line were then incubated with streptavidin-coated magnetic beads, stringently washed, and subjected to on-bead trypsin digestion. Digests of the enriched proteins were labeled with isobaric tandem mass tags (TMT) (*23*) for relative quantitation, mixed for multiplexing, and subsequently analyzed by LC-MS/MS (Figure 1B). We used a ratiometric approach of each individual sgRNA 293T-Caspex line compared to the no guide control line (*24*), which is not spatially constrained to a locus in the genome by a sgRNA. From this analysis, we detected 3,199 proteins, each with at least two quantifiable peptides. Biotinylation in four of the *hTERT* Caspex lines that according to the ChIP-qPCR results (430T, 107T, T266 and T092; Supplemental Figure 1) had overlapping labeling radii, showed high correlation of protein enrichment (Figure 2B). The T959 Caspex line, which lies approximately 2 labeling radii from its closest neighbor, showed decreased correlation of protein enrichment. We then performed a statistical analysis (moderated T-test) using each of the four overlapping sgRNA lines as quasi-replicates of each other, using the non-spatially constrained no sgRNA 293T-CasPEX line as the control. 371 proteins were significantly enriched (adj. p value < 0.05) at *hTERT* over the no sgRNA control, including five proteins known to occupy *hTERT* in various cell types (TP53; (*25*, *26*), MAZ; (*27*, *28*), CTNNB1; (*29*–*31*), ETV3; (*32*), CTBP1; (*33*) (Figure 2C, Supplemental Table 1). These results indicate GLoPro is able enrich proteins from the native, cellular context, and suggests this method is capable of distinguishing proteins at a particular genomic locus.

In addition to detecting proteins known to associate with *hTERT*, we also identified several novel candidate proteins associated with this promoter region. To corroborate whether a subset the proteins identified by GLoPro associate with *hTERT*, we performed ChIP-qPCR for several candidates. Since many of these proteins do not have ChIP grade antibodies we turned to V5-tagged ORF expression in unmodified HEK293T cells (*34*). Twenty-three individual V5-tagged ORFs were chosen by availability, having ≥ 99% amino acid homology, and having an in-frame V5 tag. We selected 16 V5-tagged ORFs that spanned a range of significant enrichment values (Figure 2D). We also chose four V5-tagged ORFs for proteins not identified as significantly enriched at *hTERT*, and three proteins that were not detected as negative controls. To moderate overexpression, each ORF was individually expressed in HEK293T cells at one-fourth of the recommended DNA concentration. After 48 hours, the cells were subjected to anti-V5 ChIP-qPCR with probes tiling the regions targeted by the sgRNAs. Comparing ChIP-qPCR signals from each individual ORF to their respective GLoPro enrichment values (proteins not detected were assigned a GLoPro enrichment value of 0) we found that all proteins enriched in the GLoPro analysis were, as a group, statistically enriched by ChIP-qPCR (Mann-Whitney test, p = 0.0008) (Figure 2E). Most candidates deemed statistically enriched according to the GLoPro analysis were separated in the ChIP enrichment space from those not enriched or not detected. Two proteins previously described to bind *hTERT*, CTBP1 and MAZ (*27*, *28*, *33*), were found in a regime of high ChIP enrichment and low GLoPro enrichment, suggesting ChIP-qPCR provides orthogonal information to GLoPro for protein occupancy at a genomic locus. These data show that GLoPro can identify known, but also novel proteins that can be corroborated by ChIP-qPCR, that associate with *hTERT.*

To explore the generalizability of GLoPro at another site in the genome, we created 293T-Caspex cells that express individual sgRNAs that tile the *c-MYC* promoter (Figure 3A). ChIP-qPCR against CASPEX verified the proper localization of each *c-MYC* 293T-Caspex line (Supplemental Figure 3). GLoPro analysis of the *c-MYC* promoter identified 66 proteins as significantly enriched (adj. p val < 0.05) compared to the no guide control line (Figure 3B, Supplemental Table 2). We applied a machine learning algorithm to identify association of GLoPro-enriched proteins with canonical pathways from the Molecular Signature Database (*35*, *36*), http://apps.broadinstitute.org/genets). We identified 21 statistically enriched networks (adj. p val. < 0.01), including the “MYC Active Pathway”, a gene set of validated targets responsible for activating *c-MYC* transcription (*36*) (Figure 3C). To corroborate the association of proteins with the c-MYC promoter identified by GLoPro, we focused on components of enriched gene sets using ChIP-qPCR. ChIP-qPCR confirmed the presence of pathway components at the c-MYC promoter, including HUWE1, RUVBL1, and ENO1 for MYC active *pathway*, RBMX for *mRNA splicing pathway*, and MAPK14 (a.k.a. P38a/MXI2) for the *Lymph_angiogenesis pathway* (Figure 3D). Taken together, these results illustrate that GLoPro enriches and identifies proteins associated in multiple pathways that are known to activate *c-MYC* expression, while directly implicating specific proteins potentially involved in regulating *c-MYC* transcription through association with its promoter.

**Figure 3.**
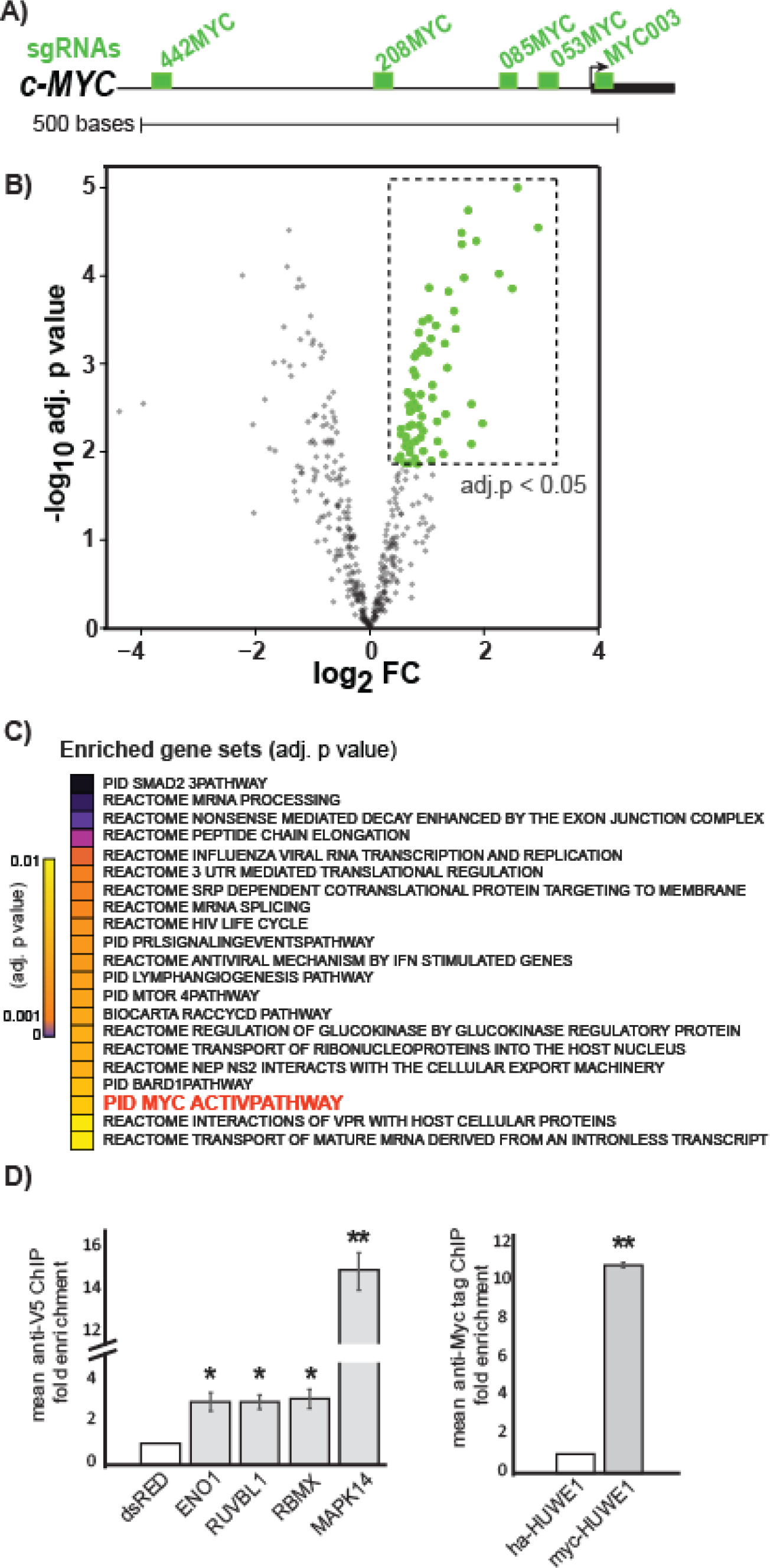
Genomic locus proteomic analysis of *c-MYC* promoter. **A)** UCSC Genome Browser representation (hg19) of the *c-MYC* promoter and the location of sgRNA sites relative to the TSS. **B)** Volcano plot of proteins quantified across the five MYC-Caspex cell lines compared to the no sgRNA control Caspex line. Data points representing proteins with an adjusted p-value of less than 0.05 are labeled green. **C)** Significantly enriched gene sets from proteins identified to associate with the *c-MYC* promoter by GLoPro. Only gene sets with an adjusted p-value of less than 0.01 are shown. MYC ACTIVE PATHWAY is highlighted in red and discussed in the text **D)** ChIP-qPCR of candidate proteins identified by GLoPro at the *c-MYC* promoter. V5 tagged dsRED served as the negative control for V5-tagged proteins ENO1, RBMX, RUVBL1 and MAPK14, whereas HA-tagged HUWE1 was used for MYC-tagged HUWE1. * indicates T-test p-value < 0.05, ** p < 0.01.

Here, we present a method for the unbiased discovery of proteins associated with particular genomic loci in live cells without genetically engineering the site of interest. We applied GLoPro to identify proteins associated with the *hTERT* and *c-MYC* promoters. Both well-established and previously unreported interactors of the respective promoter regions identified by GLoPro were validated using ChIP-qPCR, demonstrating that this method enables the discovery of proteins and pathways that potentially regulate a gene of interest without the need for prior knowledge of potential occupants.

GLoPro relies on the localization of the affinity labeling enzyme APEX2 directed by the catalytically dead CRISPR/Cas9 system to biotinylate proteins within a small labeling radius at a specific site in the genome in living cells. Other than the expression of Caspex and its associated sgRNA, no genome engineering or cell disruption is required to capture a snapshot of proteins associated with the genomic locus of interest. This advantage, in combination with the generalizability of dCAS9 and APEX2, suggests that GLoPro can be used in a wide variety of cell types and at any dCAS9-targetable genomic element. Beyond circumventing the need for antibodies for discovery, LC-MS/MS analysis using isobaric peptide labeling allows for sample multiplexing, enabling multiple sgRNA lines and/or replicates to be measured in a single experiment with little or no missing data for relative quantitation of enrichment. GLoPro-derived candidate proteins can be further validated for association with the genomic region of interest by ChIP, the current gold standard for interrogation of protein-DNA interactions. While GLoPro in this initial work only identifies association with a locus and not functional relevance, we expect that analyzing promoters or enhancer elements during relevant perturbations may provide novel functional insights into transcriptional regulation. In addition, we envision CASPEX can be used for enrichment of genomic locus entities such as locus-associated RNAs (i.e. nascent or non-coding RNAs) or DNA elements not targeted directly by CASPEX, but close in three-dimensional space within the nucleus (i.e. enhancers or promoters associated with an enhancer). Further work will be needed to assess the extended capabilities of CASPEX.

While we have demonstrated that GLoPro will be a powerful tool to study chromatin structure and transcriptional regulation, there are several drawbacks that should be noted, mainly concerning receptive cell systems and analyte sensitivity. We designed GLoPro to have an inducible expression system to prevent constant CASPEX association with the locus of interest, potentially disrupting gene expression. Thus, the inducible expression and selection cassette is currently too large for viral transduction (Figure 1C). Ongoing work in our laboratory has found that co-transfecting the piggybac transposase aids the generation of stable Caspex lines in cell culture systems with poor transfection efficiency (data not shown). Thus, in its current form, Caspex can only be used in electroporation or transfectable cells. The second major challenge is sensitivity. Avoidance of chemical crosslinking, the high affinity of streptavidin for biotin, and sample multiplexing were boons for the development of GLoPro, but due to the inherent sensitivity limits of current mass spectrometers and the unavoidable sample loss at each sample handling step, a large amount of input material is needed. These input requirements are readily attainable with many cell culture systems but may prove more challenging with recalcitrant or limited passaging cells.

In summary, we describe genomic locus proteomics, a novel approach to identify proteins associated with a predefined site in the genome. This method provides an orthogonal and highly complementary approach to ChIP for the unbiased discovery of proteins that may regulate gene expression and chromatin structure.

## Figures and Figure Legends

**Supplemental Figure 1.**
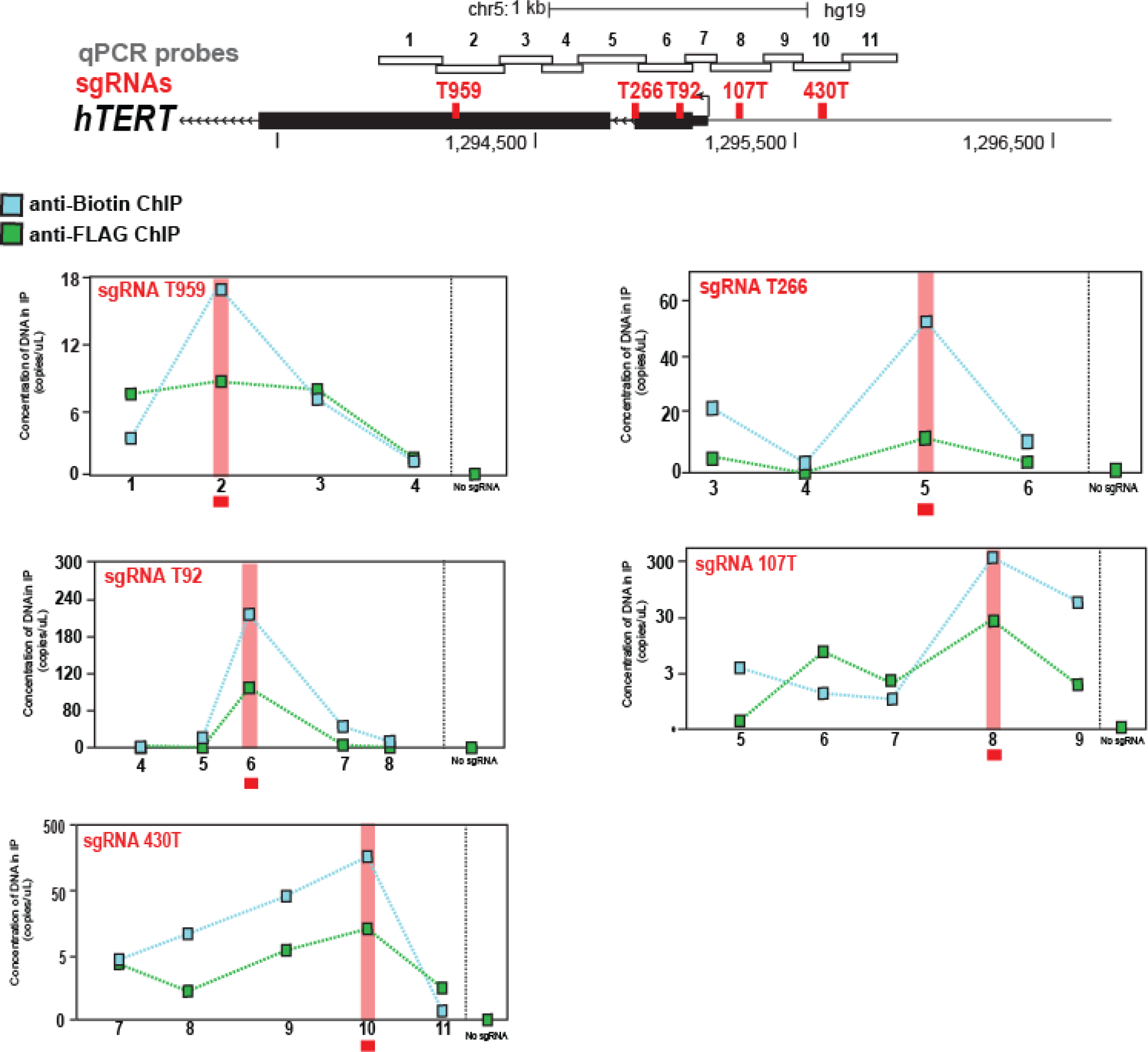
A) UCSC Genome Browser representation (hg19) of the *TERT* promoter, including genomic coordinates, and the location of sgRNA sites relative to the TSS. qPCR probes are numbered. B) ChIP-qPCR against biotin (blue boxes) and FLAG (green boxes) in hTERT-CasPEX cells expressing either no sgRNA (far right) their respective sgRNA. ChIP probes refer to regions amplified and detected by qPCR. The location of the sgRNA in each ChIP-qPCR is highlighted in red.

**Supplemental Figure 2.**
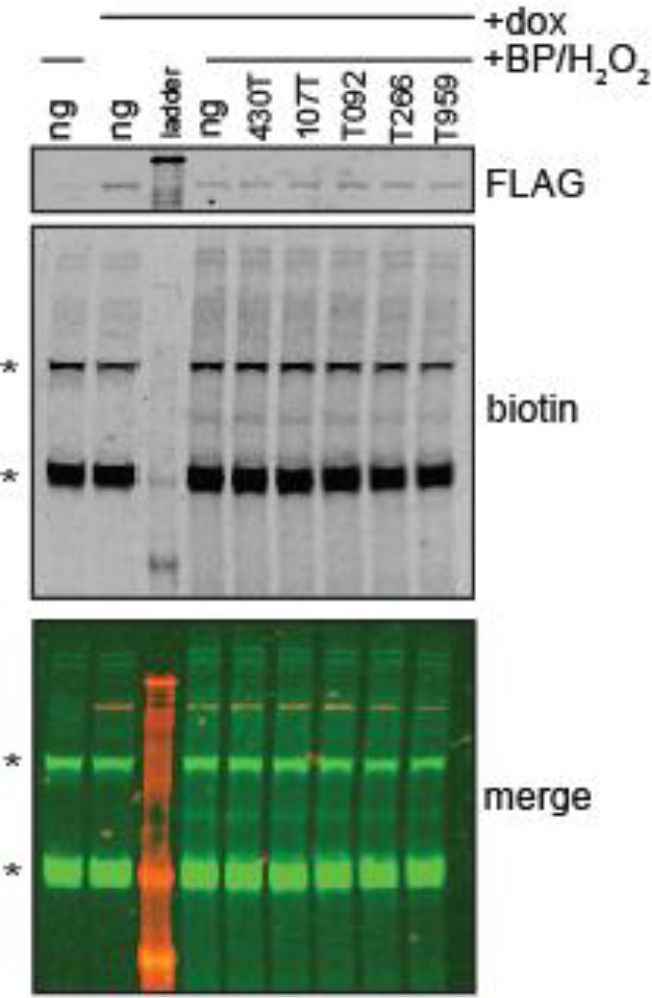
Anti-FLAG and anti-biotin Western blots of *TERT* Caspex lines treated for 12 hours with 0.5 ug/mL dox or vehicle, and labeled via Caspex-mediated biotinylation. Top panel shows anti-FLAG signals for cells treated with dox or vehicle. Middle panel shows anti-biotin signal from cells exposed to labeling protocols with or without dox treatment. Endogenous biotinylated proteins (stars) are used as the loading control. Bottom panel is a merge of both signals. Protein molecular weight ladder separates the no-guide line from the *TERT* Caspex lines.

**Supplemental Figure 3.**
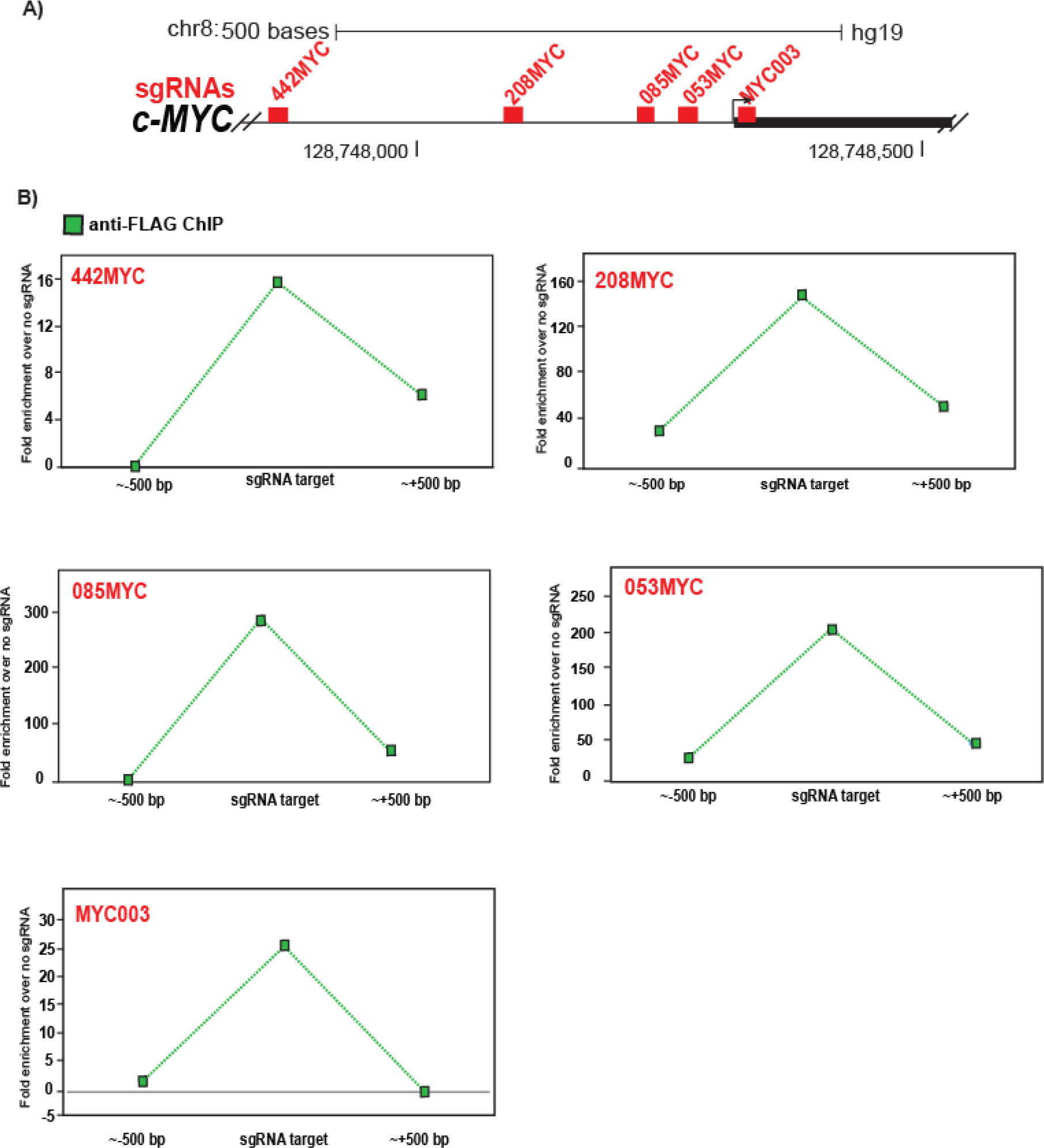
A) UCSC Genome Browser representation (hg19) of the *c-MYC* promoter, including genomic coordinates, and the location of sgRNA sites relative to the TSS. B) ChIP-qPCR against CASPEX (FLAG epitope) in MYC-Caspex cells expressing their respective sgRNAs. ChIP probes either span the region targeted by the respective sgRNA or a non-overlapping regions approximately 500 bp on either side of the sgRNA target site. Caspex cells expressing no sgRNA was used as the negative control.

**Supplemental Table 1** Values for fold enrichment of individual guides and results for the moderated T test performed on the *hTERT* GLoPro data discussed in the paper.

**Supplemental Table 2** Values for fold enrichment of individual guides and results for the moderated T test performed on the *c-MYC* GLoPro data discussed in the paper.

**Supplemental Table 3** Oligonucleotides used in this study

## Acknowledgements

We are grateful to Seth Shipman and George Church for the tetracycline-inducible/puro^R^ plasmid. We would like to thank Alice Ting, John Doench, Aviv Regev, Namrata Udeshi, and Egle Kvedaraite for useful discussion; Ryan Peckner, Elizabeth M. Perez, and Robbyn Issner for technical assistance; Malvina Papanastasiou, Eric Kuhn, Jennifer Abelin for critical assessment of the manuscript. This work is funded by NCI CPTAC PCC 1U24CA210986-01 and the Broad Institute SPARC grant. The authors will make the Caspex plasmid available to the community through Addgene. The authors declare no conflicting financial interests. A patent application has been filed by The Broad Institute relating to this work. S.A.M. and J.W. conceived the experimental set-up and designed the research. S.A.M. created the constructs and stable lines, performed the labeling, Western blots, the enrichments and the mass spectrometry, and the ChIP-qPCRs for the *c-MYC* lines. J.W. performed the ChIP-qPCRs for the *hTERT* lines. All authors interpreted the data. S.A.M., J.W. and S.A.C. wrote the manuscript.

## Methods

### Plasmid construction

The Caspex construct (dox inducible dCas9-APEX2-T2-GFP) was created by subcloning *3xFLAG-dCas9* and *T2A-Gfp* from pLV-hUBC-dCas9-VP64-T2A-GFP (Addgene 53192), and V5-APEX2-NLS from mito-V5-APEX2 (Addgene 42607) into an all in one piggybac, TREG/Tet-3G plasmid (Church lab) via ligation independent cloning (InFusion, Clontech). Guide sequences were selected and cloned as previously described (Doench *et al*). All V5 ORF constructs were purchased through the Broad Genetics Perturbation Platform and were expressed from the pLX-TRC_317 backbone. V5 ORFs were only selected for validation if the construct was available, had protein homology >98%, and an in frame V5 tag.

### Cell Line construction and culture

HEK293T cells were grown in DMEM supplemented with 10% fetal bovine serum, glutamine and non-essential amino acids (Gibco). All constructs were transfected with Lipofectamine 2000. After Caspex transfection, puroMYCin was added to a final concentration of 4 ug/ml and selected for two weeks. Single colonies were picked, expanded and tested for doxycycline inducibility of the Caspex construct monitored by GFP detection. The HEK293T cell line with the best inducibility (now referred to as 293-Caspex cells) was expanded and used for all subsequent experiments. For stable sgRNA expression, single sgRNA constructs were transfected into 293-Caspex cells and were selected for stable incorporation by hygroMYCin treatment at 200 ug/ml for two weeks. Caspex binding was tested using ChIP followed by digital droplet PCR (ddPCR) or qPCR.

### APEX-mediated labeling

Prior to labeling, doxycycline dissolved in 70% ethanol was added to cell culture media to a final concentration of either 500 ng/mL for 18-24 hours (*hTERT*) or 12 hours at 1 ug/mL (*c-MYC).* Biotin tyramide phenol (Iris Biotech) in DMSO was added directly to cell culture media, which was swirled until the precipitate dissolved, to a final concentration of 500 uM. After 30 minutes at 37°C hydrogen peroxide was added to media to a final concentration of 1 mM to induce biotinylation. After 60 seconds the media was decanted and the cells were washed with ice cold PBS containing 100 mM sodium azide, 100 mM sodium ascorbate and 50 mM TROLOX (6-hydroxy-2,5,7,8-tetramethylchroman-2-carboxylic acid) three times. Cells were lifted and transferred to 15 ml Falcon tubes with ice cold PBS, spun at 500g for 3 minutes, flash frozen in liquid nitrogen and stored at −80°C.

### Chromatin immunoprecipitation followed by quantitative PCR

Cells were trypsinized to single cell suspension and fresh formaldehyde was added to a final concentration of 1% and incubated at 37°C for 10 minutes, being inverted several times every two minutes or so. Formaldehyde was quenched with 5% glycine and the samples were aliquoted into 3e6 cell aliquots, spun down and flash frozen in 0.5 mL Axygen tubes. Chromatin was sheared using a QSonica Q800R2 Sonicator at and amplitude of 50 for 30 seconds on/30 off, for 7.5 minutes, until 60% of fragments were between 150 and 700bp. Lysis buffer was comprised of 1% SDS, 10 mM EDTA and Tris HCl, pH 8.0. For ChIP, streptavidin (SA) conjugated to magnetic beads (Thermo), M2 anti-FLAG antibody (Sigma) or anti-V5 antibodies (MBL Life Sciences) was conjugated to a 50:50 mix of Protein A: Protein G Dynabeads (Invitrogen) was incubated with sheared chromatin at 4°C overnight. qPCR was performed with either Roche 2x Sybr mix on a Lightcycler (Agilent) or via digital droplet PCR (BioRad).

### Western blot analysis

sgRNA-293-Caspex cells were labeled as described above. 40 ug of whole cell lysate was separated by SDS-PAGE, transferred to nitrocellulose and blotted against FLAG (Sigma) or biotin (Li-Cor IRdye 800 CW Streptavidin and IRdye 680RD anti-Mouse IgG).

### Enrichment of biotinylated proteins for proteomic analysis

Eight 15 cm2 plates of each sgRNA-293-Caspex line, or no guide as a negative control, were used for proteomic experiments. Labeled whole cell pellets were lysed with RIPA (50 mM TRIS pH 8.0, 150 mM NaCl, 1% NP-40 and 0.5% sodium deoxycholate, 0.1% sodium dodecyl sulfate) with protease inhibitors (Roche) and probe sonicated to shear genomic DNA. Whole cell lysates were clarified by centrifugation at 14,000g for 30 minutes at 4°C and protein concentration was determined by Bradford. 500 uL SA magnetic bead slurry (Thermo) was used for each sgRNA line (between 60-90 mgs of protein/state). Lysates of equal protein concentrations were incubated with SA for 120 minutes at room temperature, washed twice with cold lysis buffer, once with cold 1M KCl, once with cold 100 mM Na_3_CO_2_, and twice with cold 2 M urea in 50 mM ammonium bicarbonate (ABC). Beads were resuspended in 50 mM ABC and 300 ng trypsin and digested at 37oC overnight.

### Isobaric labeling and liquid chromatography tandem mass spectrometry

On-bead digests were desalted via Stage tip (*37*) and labeled with TMT (Thermo) using an on-column protocol. For on-column TMT labeling, Stage tips were packed with one punch C18 mesh (Empore), washed with 50 uL methanol, 50 uL 50% acetonitrile (ACN)/0.1% formic acid (FA), and equilibrated with 75 uL 0.1% FA twice. The digest was loaded by spinning at 3,500g until the entire digest passed through. The bound peptides were washed twice with 75 uL 0.1% FA. One uL of TMT reagent in 100% ACN was added to100 uL freshly made HEPES, pH 8, and passed over the C18 resin at 2,000g for 2 minutes. The HEPES and residual TMT was washed away with 75 uL 0.1% FA twice and peptides were eluted with 50 uL 50% ACN/0.1% FA followed by a second elution with 50% ACN/20 mM ammonium hydroxide, pH 10. Peptide concentrations were estimated using an absorbance reading at 280 nm and mixed at equal ratios. Mixed TMT labeled peptides were step fractionated by basic reverse phase on a sulfonated divinylbenzene (SDB-RPS, Empore) packed Stage tip into 6 fractions (5, 10, 15, 20, 30 and 55% ACN in 20 mM ammonium hydroxide, pH 10). Each fraction was dried via vacuum centrifugation and resuspended in 0.1% formic acid for subsequent LC-MS/MS analysis.

Chromatography was performed using a Proxeon UHPLC at a flow rate of 200 nl/min. Peptides were separated at 50°C using a 75 micron i.d. PicoFit (New Objective) column packed with 1.9 um AQ-C18 material (Dr. Maisch) to 20 cm in length over a 94 min gradient. Mass spectrometry was performed on a Thermo Scientific Q Exactive Plus (*hTERT* data) or a Lumos (*c-MYC* data) mass spectrometer. After a precursor scan from 300 to 2,000 m/z at 70,000 resolution the top 12 most intense multiply charged precursors were selected for HCD at a resolution of 35,000. Data were searched with Spectrum Mill (Agilent) using the Uniprot Human database, in which the CASPEX protein was amended. A fixed modification of carbamidomethylation of cysteine and variable modifications of *N*-terminal protein acetylation, oxidation of methionine, and TMT-10plex labels were searched. The enzyme specificity was set to trypsin and a maximum of three missed cleavages was used for searching. The maximum precursor-ion charge state was set to 6. The precursor mass tolerance and MS/MS tolerance were set to 20 ppm. The peptide and protein false discovery rates were set to 0.01.

### Data analysis

All non-human proteins and human proteins identified with only one peptide were excluded from downstream analyses. Human keratins were included in all analyses but were removed from the figures. The moderated T-test was used to determine proteins statistically enriched in the sgRNA-293-Caspex lines compared to the no sgRNA control. After correcting for multiple comparisons (Benjamini-Hochberg procedure), any proteins with an adjusted p-value of less than 0.05 were considered statistically enriched.

Pathway analysis was performed using the Quack algorithm incorporated into Genets (http://apps.broadinstitute.org/genets) to test for enrichment of canonical pathways in the Molecular Signature Database (MSgiDB). Proteins identified as significantly enriched (adj. p-val. < 0.05) by GLoPro were input into Genets and were queried against MSigDB. Pathways enriched (FDR < 0.05) were investigated manually for specific proteins for follow-up.

